# Ψ-Atlas: an integrated atlas for pseudouridine epitranscriptome

**DOI:** 10.1101/2025.06.06.656536

**Authors:** Xiaochen Wang, Jinjing Luo, Xiaoqiang Lang, Yongqing Ling, Yiming Zhou, Guoxian Liu, Xiangye Chen, Yibo Chen, Yingshun Zhou, Yi Cao, Zhonghui Zhang, Changjun Ding, Demeng Chen, Qi Liu

## Abstract

Pseudouridine (Ψ) is a C^5^ glycoside isomer of uridine, formed by breaking the N^1^ glycosyl bond and undergoing a 180° base rotation. This modification stands as one of the most widespread post-transcriptional alterations in RNA and is universally distributed among diverse RNA species. The pervasiveness of this modification enhances RNA structural integrity, bestows unique structural and functional attributes upon the RNA molecules it adorns, and facilitates additional hydrogen bonding. However, the absence of a convenient and integrated intuitive visualization database encompassing all currently reported species and RNA types is evident. Here, we present the Ψ-Atlas, an extensive database meticulously curated for the comprehensive collection and annotation of RNA pseudouridine. This database encompasses 554,895 pseudouridine modification sites across various RNA categories, including mRNA, ncRNA, tRNA, rRNA, and others, in 55 distinct species reported in current literature. The sequencing methodologies employed comprise the next-generation sequencing (NGS) techniques such as Ψ-Seq, Pseudo-Seq, CeU-Seq, PSI-Seq, RBS-Seq, HydraPsiSeq, BID-Seq, PRAISE-Seq, as well as third-generation sequencing methods like Direct RNA sequencing (DRS). Ψ-Atlas as the most comprehensive and integrated resource for RNA pseudouridine modifications to date. And the Ψ-Atlas database offers an intuitive interface for information display and a myriad of analytical tools, including PsiVar and PsiFinder. In essence, this platform serves as a robust search and visualization tool for the study of pseudouridylation in epitranscriptomics. Ψ-Atlas is available at https://rnainformatics.org.cn/PsiAtlas.

## Introduction

Currently, over 170 distinct ribonucleotide modifications occurring on RNA have been identified [1,2]. Among these, pseudouridine, an isomer of uridine (U) at the C5-glycoside, stands out as one of the earliest identified and most abundant ribonucleotide modifications [3]. It is ubiquitously present in both coding RNA and various non-coding RNA types, including rRNA, tRNA, long non-coding RNA (lncRNA), small nuclear RNA (snRNA), and small nucleolar RNA (snoRNA) [3–6]. Many Ψ positions in rRNA are highly conserved. For instance, Escherichia coli 23S rRNA includes three conserved Ψ residues at positions 1911, 1915, and 1917 [6,7]. Mitochondrial rRNA is encoded by the mitochondrial genome and is subject to pseudouridylation. One identified modification site is Ψ1397 in *Homo sapiens* mitochondrial LSU 16S rRNA, located near the peptidyl transferase center (PTC) core [8]. In most tRNAs, Ψ is found not only in the conserved TΨC loop but also in the D stem and/or anticodon stem and loop [9]. Additionally, Ψ is conservatively present in spliceosomal snRNAs, such as U1, U2, U4, U5, and U6. Its specific locations are primarily within regions crucial for RNA-RNA and RNA-protein interactions essential for spliceosome assembly and splicing processes [10–12]. While in the early stages of Ψ research, non-coding RNA was the primary focus. However, with the advancement of high-throughput technologies, an increasing number of Ψ sites have been discovered on mRNA in *Saccharomyces cerevisiae* and *Homo sapiens* cells [13–15]. Early investigations into the structure of Ψ residues have, to a certain extent, laid the groundwork for later elucidating the functional implications of pseudouridylation occurring on RNA. Due to the presence of two hydrogen donors (N^1^ and N^3^ imino protons) and two hydrogen acceptors in Ψ [16], when Ψ forms pairs within the double helix with A, G, U, or C, creating Ψ-A, Ψ-G, Ψ-U, and Ψ-C pairs, it enhances the thermodynamic stability of RNA duplexes. This results in more stable base stacking arrangement and a more rigid RNA structure compared to uridine [16–18]. These structures and thermodynamic properties contribute to the specific functions elicited by Ψ in vivo. For instance, pseudouridylation of rRNA optimizes the function of distinct ribosomal subpopulations, enabling the translation process and cellular proteome to adapt to specific intracellular or environmental conditions [19]. Depletion of the *RluD* gene in *E. coli* results in the absence of Ψ at positions 1911, 1915, and 1917 in the 23S rRNA, leading to a significant inhibition of *E. coli* growth [20]. Depletion of the essential enzyme RNA Pseudouridine Synthase D4 (RPUSD4), responsible for the Ψ1397 modification in *Homo sapiens* mitochondrial LSU 16S rRNA, reduces the activity of electron transport chain complexes, thereby impeding mitochondrial translation [8]. In *A. thaliana*, Leaf Curly and Small 1 (FCS1), located in the mitochondria, has been identified as a mediator of pseudouridylation in mitochondrial 26S rRNA. In *fcs1* mutants, although mitochondrial gene transcripts increase, protein accumulation is reduced. This results in severe impairment of mitochondrial structure and function, consequently affecting the growth and development of *A. thaliana* [4]. In tRNA, Ψ stabilizes its secondary structure, influencing the rate and accuracy of translation, thereby playing a pivotal role in directing stem cell translational control [21]. Ψ residues in snRNA aggregate in crucial functional regions, actively participating in precursor mRNA splicing. On spliceosomal U6 snRNA, the Ψ residue at position 28 (Ψ28), catalyzed by Pseudouridine Synthase 1p (Pus1p) during *Saccharomyces cerevisiae* filamentous growth, contributes to this filamentous growth program [22]. Pseudouridylation contributes to the development of COVID-19 mRNA vaccines by reducing the immunogenicity of mRNA vaccines, thereby deceiving the immune system [23–25].

The synthesis of Ψ in vivo is generally mediated by enzymes, known as pseudouridine synthase (PUS). Within this synthase superfamily, members of the tRNA pseudouridine(38-40) synthase (TruA), tRNA pseudouridine(55) synthase (TruB), and tRNA pseudouridine(13) synthase (TruD) families are involved in the modification of tRNA, while members of the bifunctional tRNA pseudouridine(32) synthase/23S rRNA pseudouridine(746) synthase (RluA) and RsuA families primarily participate in pseudouridine modifications of rRNA [26]. This synthesis mechanism is universally present across diverse organisms, including eukaryotes, prokaryotes and archaea [27]. For instance, TruB (bacterial origin) and Pus10 (archaeal origin) have demonstrated the ability to modify the U at position 55 of tRNA to Ψ [26,28,29]. Another mechanism for the formation of Ψ involves the catalysis by H/ACA box ribonucleoproteins (RNPs) containing the ANANNA or ACA motifs, leading to the isomerization of specific Us in rRNA, U2 snRNA, and mRNA into Ψ [3,30].

In addition to the early identification of Ψ using mass spectrometry (MS) [31], detection techniques for Ψ have continually evolved with the advancement of sequencing technologies. Specifically, chemical sequencing methods derived from NGS technologies, such as Ψ-Seq, Pseudo-Seq, PSI-Seq, CeU-Seq, and others, have been developed [32–35]. Recently, with the introduction of DRS, the mapping of Ψ has reached a new pinnacle [36]. Therefore, the integration of a comprehensive Ψ map database covering all species and RNA types becomes crucial. However, there are only a few comprehensive databases that contain a small number of pseudouridine modification sites, such as RMDisease V2.0 [37], DirectRMDB [36], RMBase v3.0 [38] and PRMD [39], and to date, there is no one dedicated Ψ modification database spanning multi-species, multi-technology databases. In this regard, we have established the Ψ-Atlas website, encompassing 55 species and 554,895 Ψ modification sites (https://rnainformatics.org.cn/PsiAtlas). This platform serves as a robust search and visualization tool for the study of pseudouridylation in epitranscriptomics.

## Results

### **Ψ** sites collected in **Ψ**-Atlas

We developed Ψ-Atlas as the most comprehensive and integrated resource for RNA pseudouridine modifications to date (Figure 1). Ψ-Atlas spans 55 species, comprising 20 animals, 15 plants, and 43 microorganisms, and the database contains 554,895 Ψ sites, collected from studies utilizing NGS and DRS technologies, and we additionally gathered all published pseudouridine modification data obtained through other experimental methods, such as MS and antibody-based sequencing, along with corresponding RNA types and species information, including *Homo sapiens* (301,335), *Mus musculus* (113,961), *Saccharomyces cerevisiae* (12,270), *Arabidopsis thaliana* (10,509), *Cucumis melo* (10,683), *Penaeus monodon* (20,364), and others species (Figure 2A and Supplementary Table S3). Moreover, Ψ-Atlas collected data on the number of pseudouridine modifications, RNA types, and other information across 16 disease types in human, including Acute Lymphoblastic Leukemia, Cervical Carcinoma, Diabetes, and Dyskeratosis Congenita (Figure 2B). The related data is available for download on the Download page. All the datasets mentioned above were processed through our standardized pipeline, with information stored in a MySQL database and displayed in a user-friendly web module within Ψ-Atlas. Furthermore, we developed convenient analytic tools that allow users to interactively analyze uploaded datasets, such as PsiFinder for enabling the identification of putative Ψ sites in genomic regions of interest and PsiVar for detecting the potential effects of genetic variants on pseudouridine modification sites.

**Figure 1.**
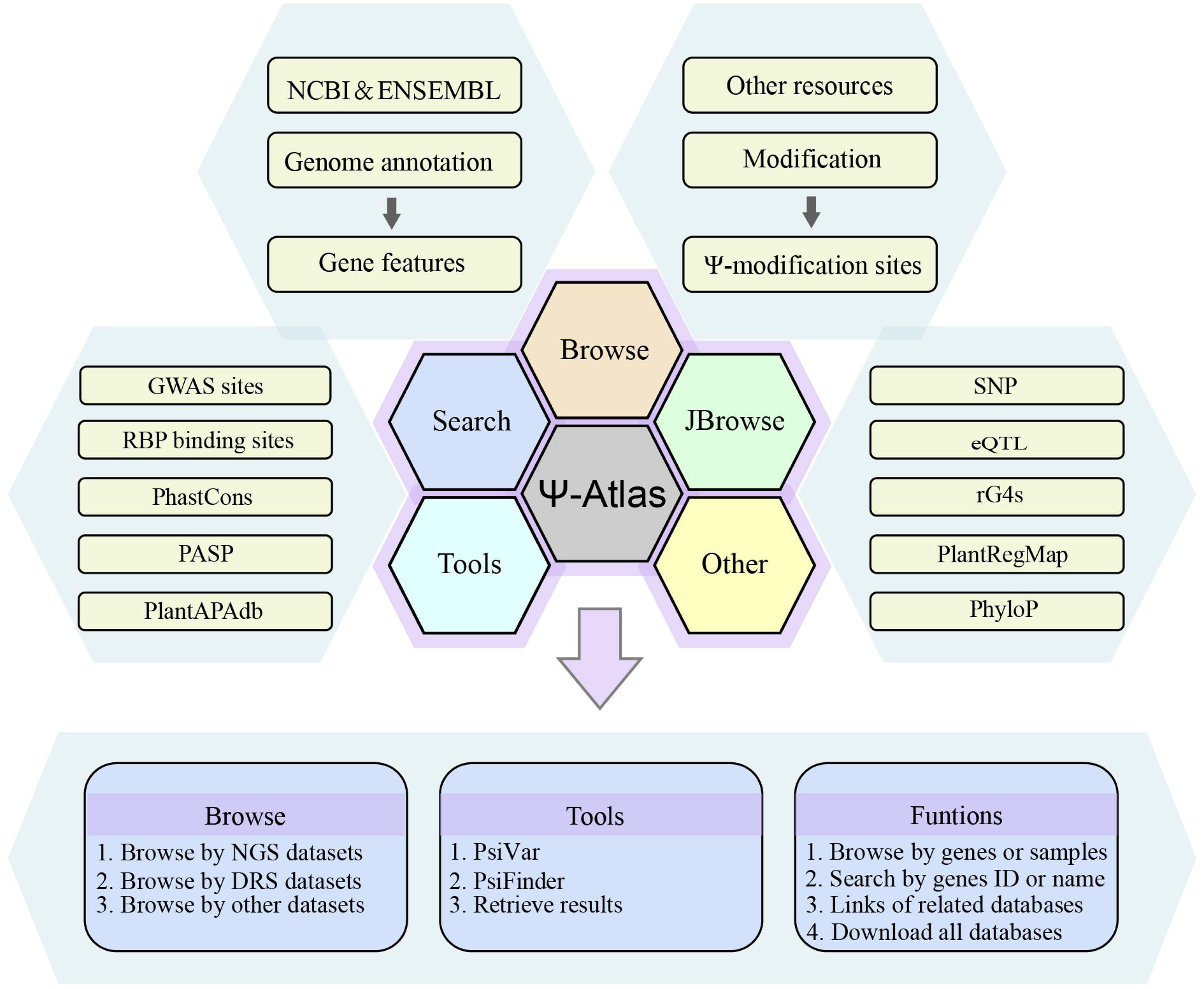
Overall workflow of. Ψ**-Atlas.** Ψ-Atlas is a comprehensive and extensive database that systematically collects and annotates Ψ modification sites. The database contains 554,895 Ψ modification sites reported in the current literature from 55 different species, including mRNA, ncRNA, tRNA, rRNA, and other RNA categories. Ψ-Atlas offers an intuitive interface for information display and two analytical tools, including PsiVar and PsiFinder.

**Figure 2.**
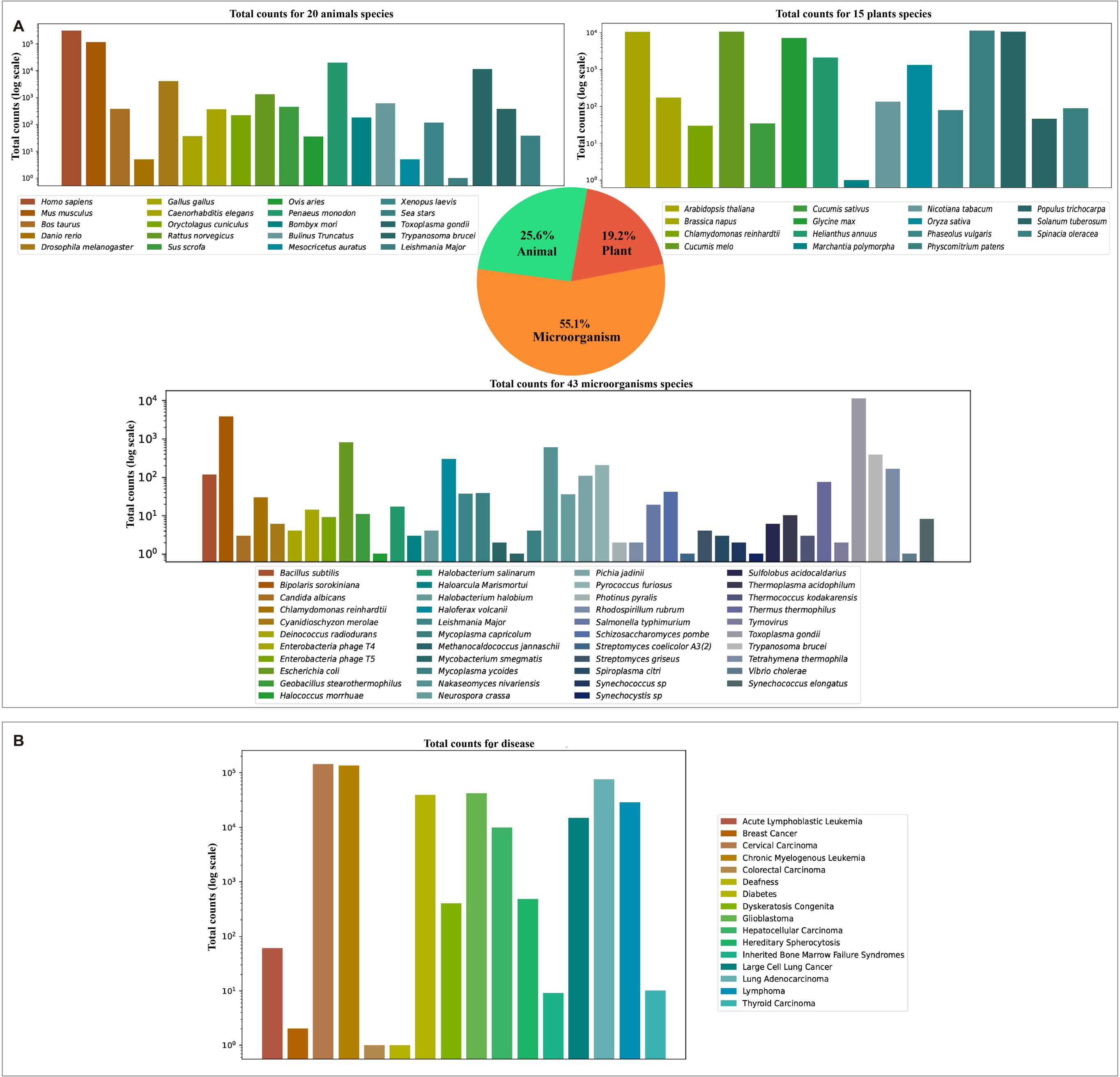
Ψ **sites in** Ψ**-Atlas across species and diseases.** (A) Statistics of the Ψ sites for each species. (B) Statistics of Ψ sites for human disease.

### Collection of **Ψ** sites based on profiling techniques

The methods encompassed in this database for detecting Ψ sites include mass spectrometry-based or chromatography-based technique, such as LC-MS [31,40–42], SILNAS [43], pseudo-MS^3^ [44], 2D-TLC [45]. As well as chemical-based or antibody-based approach coupled with NGS technologies, exemplified by PRAISE [46], Ψ-Seq [47], Pseudo-Seq [34], CeU-Seq [32], PSI-Seq [35], small RNA Ψ-Seq [48], RBS-Seq [49], HydraPsiSeq [14], BID-Seq [50], CLAP (not coupled with high-throughput sequencing)[51], PA-Ψ-Seq[52]. It also involves the subsequent emergence of “third-generation” sequencing technologies based on nanopores, such as nanoRMS [53], NanoPsu [54], DRS [36].

### Enhanced web interface and usage in **Ψ**-Atlas

The database interface of Ψ-Atlas has been meticulously designed to provide a comprehensive, rapid, and user-friendly centralized knowledge repository for Ψ sites research, allowing users to swiftly query, customize searches, and freely download all gathered datasets. Four primary modules are introduced in Ψ-Atlas, namely ‘Browse’, ‘Search’, ‘JBrowse’, and ‘Tools’.

The ‘Browse’ module is divided into three main dataset groups: (i) Browse of NGS dataset; (ii) Browse of DRS dataset; and (iii) Browse of other dataset, allowing users to easily distinguish between different sources of Ψ modification data. Both the ‘Browse of NGS dataset’ and ‘Browse of DRS dataset’ groups can be filtered by species, tissue/cell line, RNA type, and genomic region. In addition to basic information for each Ψ site (e.g., genome version, gene ID, and gene name), users can obtain detailed genomic location and sequence information through the ‘More’ option (Figure 3). The ‘Browse of NGS dataset’ group includes a total of 41 species, comprising 12 animals, 9 plants, and 20 microorganisms, while the ‘Browse of DRS dataset’ group includes 18 species, comprising 6 animals, 5 plants, and 7 microorganisms. The ‘Browse of other dataset’ section includes two data tables: the Ψ sites validated by other methods and the other related Ψ sites research. The other datasets contain Ψ sites identified through experimental methods such as liquid chromatography-tandem mass spectrometry (LC-MS/MS) and two-dimensional thin-layer chromatography (2D-TLC).

**Figure 3.**
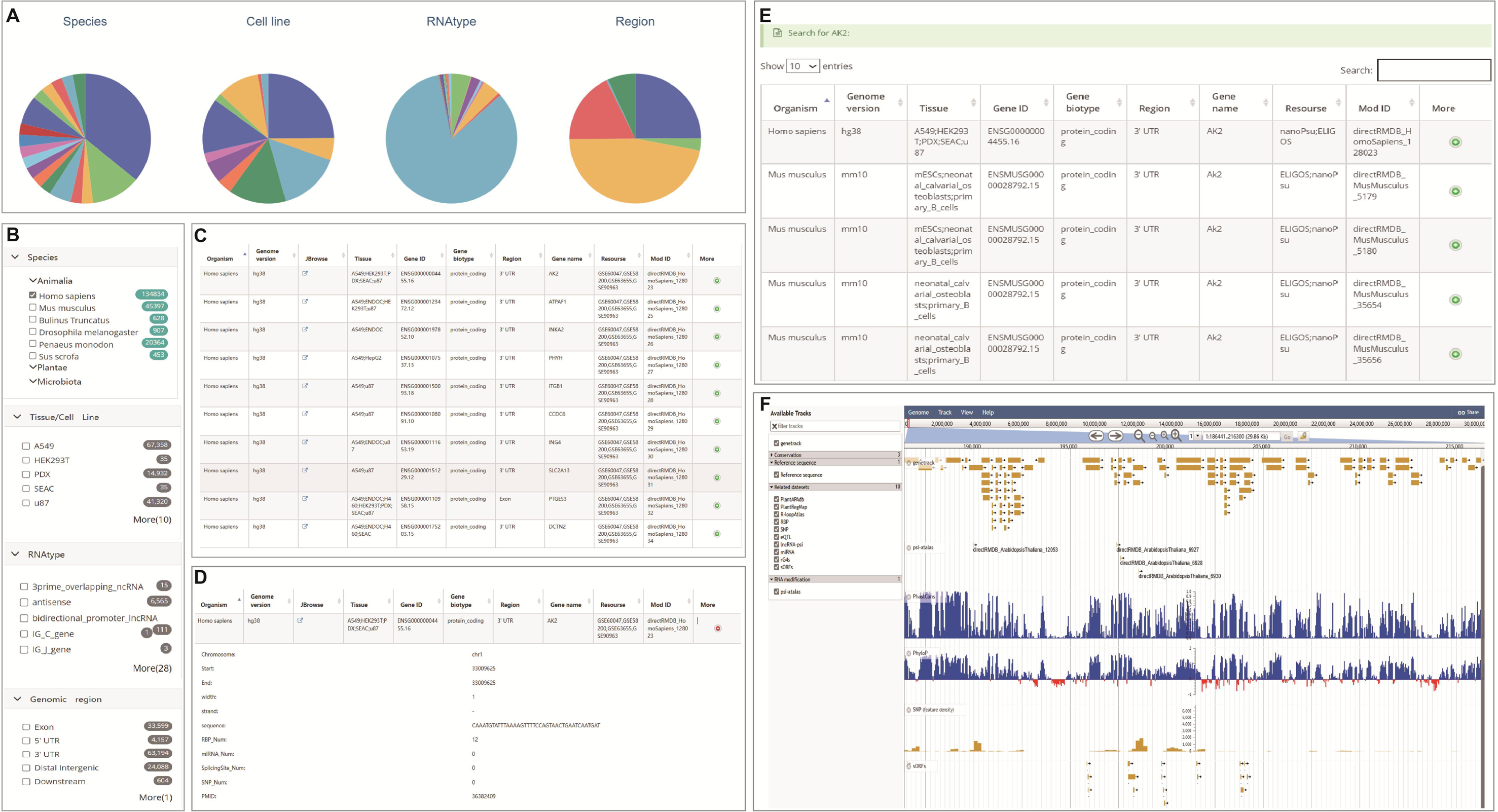
The Browse module: the collected datasets are divided into four groups based on species, tissue, RNA type, and genomic region. (A) Users can view the data distribution through pie charts. (B) Users can select categories of interest to update the table and query the data. (C) The table displays basic information of the Ψ sites. (D) Clicking ‘More’ to obtain detailed information. (E) The results of ‘Search’ module. (F) Visualization of JBrowse data displaying tracks of all Ψ modification sites and other publicly available genome annotations.

The ‘Search’ module allows users to quickly retrieve comprehensive information on Ψ modifications by submitting gene IDs, gene names, or PubMed IDs. Moreover, quickly search function is available on the Ψ-Atlas homepage for single-query searches (Figure 3E).

The ‘JBrowse’ module is integrated into a swift and scalable genome browser constructed with JavaScript and HTML5, enabling visualization of all Ψ modification sites and other publicly available genome annotations, such as RBP binding sites, miRNA targets, splice sites, SNVs, and SNPs (Figure 3F).

The ‘Tools’ module consists of following three sections: (a) the ‘PsiVar’ is used to identify the potential variant effects of pseudouridine modifications; (b) the ‘PsiFinder’ is a high-precision pseudouridine predictor based on deep learning, capable of identifying pseudouridine across various species; (c) the ‘Retrieve results’ allows users to quickly check the status of their submitted analysis tasks by job ID. The other modules include: (i) the ‘Download’ module, which provides datasets for various sequencing methods and species, enabling users to download data by ‘Species’ and ‘Data type’, as well as information related to Ψ modifications in human diseases; (ii) the ‘Links’ module, which offers links to numerous software tools and databases for Ψ modifications; and (iii) the ‘Help’ module, which provides a series of tutorials and annotations through demonstrations.

### Web-based analytical tools developed in **Ψ**-Atlas

PsiVar aims to detect potential variants affecting Ψ modification. Variant files in VCF format serve as inputs for PsiVar. Upon data submission, the queuing system assigns a job ID that users can use to monitor analysis progress in current analysis page or retrieve results of the job in the tools module. In addition, users have the option to receive email notifications upon job completion (Figure 4A). The results include a pie chart showing the distribution of Ψ loss, Ψ gain, and unchanged variants, as well as a table with detailed information, including variant position, transcript ID, peak ID, reference motif, mutated motif, and scores used to determine Ψ loss or gain. For example, PsiVar analyzed the first 10,000 rows sequence characteristics of sites with mutated variants using the SNP data of *A. thaliana* from the EnsemblPlants database (https://ftp.ensemblgenomes.ebi.ac.uk/pub/plants/release-60/variation/vcf/arabidopsis_thaliana/). Approximately 1.08% of mutated sites were classified as Ψ loss variants, while 0.29% were classified as Ψ gain variants (Figure 4B and Supplementary Table S4).

**Figure 4.**
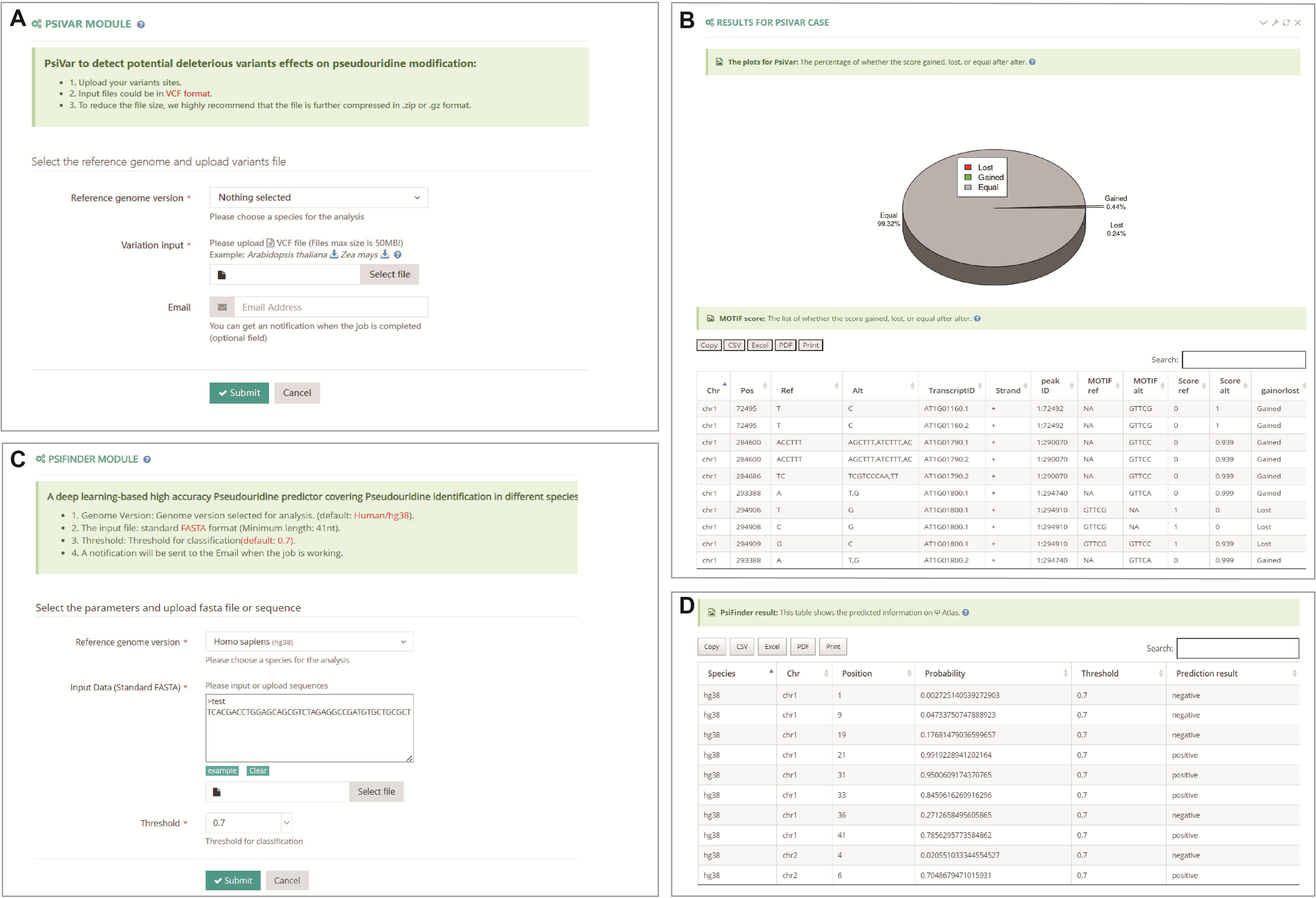
Web-based analytical tools. (A) The PsiVar is used to identify the potential variant effects of Ψ modifications. (B) The PsiFinder is a high-precision Ψ predictor based on deep learning, capable of identifying Ψ across various species. (C) The results of the score gained, lost, or equal after alter in PsiVar. (D) The predicted results of PsiFinder in Ψ-Atlas.

PsiFinder was developed to accurately predict putative Ψ sites from user-uploaded sequences in standard FASTA format. The uploaded sequences are one-hot encoded and subsequently fed into a pre-trained model for prediction. The input data is processed through two convolutional layers, followed by a normalization layer and a ReLU activation function, before being mapped to the final output via a pooling layer and a fully connected layer (Figure 4C). As an example, users can upload a randomly selected 41nt FASTA sequence from human, specify the corresponding species and set a prediction threshold (defaulting to 0.7). The predictions are then displayed in a table with fields such as Species, Chromosome, Position, Probability, Threshold, and Prediction Result. Based on these results, users can evaluate the likelihood of potential Ψ modification sites (Figure 4D).

## Discussion

Currently, several databases and web servers integrate existing pseudouridine modification sites, however, these databases are not specifically designed for Ψ modifications and cover a limited number of species and Ψ sites (Table 1). For example, RMDisease V2.0 [37] identified a total of 1,366,252 RNA modification (RM)-associated variants that may affect 16 different types of RNA modification in 20 organisms while only four species (human, mouse, yeast and arabidopsis) contained a small number of identified pseudoguanosine modification sites. DirectRMDB [36] is an Oxford Nanopore Technologies (ONT) based database that includes 16 types of modification and a total of 904,712 modification sites in 25 species while only contains DRS datasets and there has no experimental Ψ sites and NGS Ψ sites in the database. EPITOMY (https://epitomy.luddy.indianapolis.iu.edu) only focus on human and mouse and with limited pseudouridine researches. PRMD [39] is a very convenient and comprehensive database of plant RNA modifications while the analysis modules only focused on plant species as well as the most m^6^A modifications, with few studies on pseudouridine modifications. RMBase v3.0 [38] is a comprehensive and convenient platform for efficient studying RNA modifications from large amounts of epitranscriptome high-throughput sequencing data, it provides multiple interfaces and web-based tools to integrate 73 types of RNA modification among 62 species while pseudouridine modifications studies were not the main content of the database. In summary, none of these databases was developed specifically for pseudouridine modification research and based on the urgent need for a database focused on the integration of pseudouridine modification dataset, we developed Ψ-Atlas by collecting and processing all previous published pseudouridine modification data on 55 species. Additionally, Ψ-Atlas incorporated convenient analysis modules such as PsiVar and PsiFinder for data analyzes as well as JBrowse embedded for data visualization. Moreover, Ψ-Atlas intuitively displays the related datasets from previous studies and other resources such as SNVs, GWAS, RNA secondary structures, rG4 structures and RBP binding sites.

## Conclusion

Among over 170 identified ribonucleotide modifications, pseudouridine is one of the most abundant [1]. Several studies have demonstrated that Ψ modifications in RNA influence growth, development, and both physiological and pathological processes across diverse species [4,8,20]. Although several published databases have integrated existing sequencing datasets and Ψ-related information, a comprehensive and user-friendly database specifically dedicated to Ψ modification is still lacking. Therefore, we developed Ψ-Atlas, spanning 55 species, to facilitate research on Ψ modifications. Currently, Ψ-Atlas integrates 554,895 Ψ modification sites from both NGS, DRS technologies and other experimental chemical technologies. With continuous advancements in high-throughput sequencing technologies, the availability of Ψ-related sequencing data is rapidly increasing. To address this, we will regularly collect newly published high-throughput sequencing data to enrich the Ψ-Atlas database, providing more comprehensive information on Ψ modifications. Besides, we plan to develop more tools for functional analysis of Ψ-Atlas datasets, facilitating research and progress in this field.

## Methods

### Collection of **Ψ** modifications datasets

We collected and integrated all Ψ modification data currently published from 132 studies, including databases, sequencing datasets, and experimental data validated by non-high-throughput methods such as liquid chromatography-mass spectrometry (LC-MS) (Supplementary Table S1). Furthermore, we compiled the accession numbers of all raw data sources in Supplementary Table S1. For data with precise genomic locations, the sources primarily include chemical methods combined with NGS technologies, such as Ψ-Seq [47], CeU-Seq [32], and PSI-Seq [35], as well as high-throughput sequencing data obtained through nanopore-based “third-generation” sequencing technologies, including nanoRMS [53], NanoPsu [54], and DRS [36]. We integrated Ψ modification sites validated by wet-lab experiments but lacking genomic coordinates into a unified dataset under the ‘Browse of other dataset’ section of Ψ-Atlas. Using published methods, we assembled comprehensive Ψ modification information covering 55 species.

### Genome sequences of 55 species and other related datasets

The reference genome files and the corresponding genome annotation files were downloaded from GENCODE [55] for human and mouse, Ensembl and NCBI for other species. The program Liftover (https://genome-store.ucsc.edu/) was used to perform the conversion between different versions of the reference genome and the software Gffread [56] was used to convert the GFF3 and GTF file formats to each other. We also integrated additional related datasets which had intersecting factors with the Ψ modifications, including the common RBP binding datasets from the POSTAR3 database [57], genome-wide association study (GWAS) datasets from the GWAS Atlas database [58], small open reading frame (sORF) datasets from the PsORF database (http://psorf.whu.edu.cn), expression quantitative trait locus (eQTL) datasets from the AtMAD [59] and Rice-eQTL (http://riceqtl.ncpgr.cn/) databases, RNA loop information datasets from the R-loopAtlas database [60], APA site datasets from the PlantAPAdb database [61], conservation datasets from the PlantRegMap database [62], and the RNA structural information datasets such as G-quadruplex (rG4) from the G4Atlas database [63] and transcriptome-scale RNA secondary structure probing datasets from the RNA Atlas of Structure Probing (RASP) database [64] (Supplementary Tables S2).

### Prediction and functional annotation for **Ψ** sites

In addition to collecting Ψ modification data from extensive literature and databases, we obtained direct RNA nanopore sequencing samples (FAST5 data) from *Arabidopsis thaliana* (ENA: PRJEB53881), *Glycine max* (ENA: PRJEB44525), *Oryza sativa* (NCBI: PRJNA943203), *Populus trichocarpa* (NCBI: PRINA672182), and *Zea mays* (NCBI: PRJNA635654) through the NCBI and ENA databases. Firstly, the raw FAST5 signal data were converted into FASTQ format using Guppy (v6.0.1 available at https://community.nanoporetech.com/), followed by analysis with the NanoPsu software to predict pseudouridine modification sites. To ensure accuracy, a probability threshold of 0.9 was applied to filter high-confidence modification sites, resulting in a reliable dataset of pseudouridine sites across different plant species. For the prediction files of NanoPsu, further annotation was executed with some precisely modified scripts from our own RNAmod process pipeline [65]. The annotation results were subsequently displayed on the browse by DRS page of our Ψ-atlas database.

### Analysis tools developed in **Ψ**-atlas database

PsiVar was designed to detect the potential affecting of genetic variants on pseudouridine modifications. The input file must be a standard VCF file and the uploaded SNPs will be analyzed to assess their impact on the collected Ψ sites in specific species. After analyzing the input sequence characteristics, PsiVar model will give a determination of variant sites, including Ψ-gained, Ψ-lost and equal.

PsiFinder, a deep learning-based tool designed to enhance the identification and analysis of Ψ modification sites. PsiFinder employs a convolutional neural network (CNN) model, trained and tested on datasets derived from human and mouse transcriptomes. The dataset is randomly partitioned into training, validation, and test sets, comprising 80%, 10%, and 10% of the data, respectively, with the training set consisting of 14,520 samples. For each target site (T-base site) to be predicted, a 41-nucleotide window is used, which includes 20 nucleotides upstream and 20 nucleotides downstream of the site as input data. Transcripts containing Psi-Seq validated Ψ-modification sites are designated as positive samples, while randomly selected T-base site, along with their surrounding 20-nucleotide sequences, are used as negative samples. To construct a balanced dataset, equal numbers of positive and negative samples are randomly selected from the total pool of 41nt sequences. A deep neural network is trained on each individual dataset, and an ensemble model of binary classifiers is created. The final prediction score for a given input sequence is determined by averaging the prediction scores from all classifiers in the ensemble.

### Database and web interface implementation

The Ψ-Atlas database is a platform developed using HTML, PHP, CSS and Javascript, hosted in an Apache environment on a Linux system equipped with four Octa-core AMD processors (each clocked at 2.6 GHz) and 512 GB of RAM. All collected and processed datasets are stored in a MySQL database system, with interactive visualization and querying based on JQuery and DataTables. JBrowse [66] is utilized as a web-based genome browser to display the distribution of modification site on the genomic scale.

## Data availability

All previously published public data used in this study, including those from the Gene Expression Omnibus and NCBI SRA databases, are presented in Supplementary Table S1. Integrated and reprocessed data from this database are available in the ‘Browse of other dataset’ and ‘Download’ modules of Ψ-Atlas.

Ψ-Atlas is freely available at https://rnainformatics.org.cn/PsiAtlas.

## CRediT author statement

**Xiaochen Wang:** Investigation, Methodology, Data curation, Writing – original draft, Writing – review & editing. **Jinjing Luo:** Methodology, Data curation, Formal analysis, Software, Writing – original draft. **Xiaoqiang Lang:** Investigation, Methodology, Data curation, Formal analysis, Writing – original draft. **Yongqing Ling:** Software, Formal analysis. **Yiming Zhou:** Data curation, Formal analysis. **Guoxian Liu:** Investigation, Data curation. **Xiangye Chen:** Investigation, Data curation. **Yibo Chen:** Writing – review & editing, Supervision. **Yingshun Zhou:** Writing – review & editing, Supervision. **Yi Cao:** Writing – review & editing. **Zhonghui Zhang:** Writing – review & editing. **Changjun Ding:** Supervision, Funding acquisition. **Demeng Chen:** Methodology, Supervision, Funding acquisition. **Qi Liu:** Conceptualization, Methodology, Project administration, Resources, Supervision, Funding acquisition, Writing – review & editing.

## Acknowledgements

We thank the researchers from the Chen Lab for their valuable discussions and support in data analysis. This study was funded by the Basic Research Fund of CAF (CAFYBB2023QB003), National Key Research and Development Program of China (2021YFD2201205), Guangzhou Key Research and Development Program (2024B03J1319), Guangdong Basic and Applied Basic Research Foundation (2020B1515020007), Project of Collaborative Innovation Center of GDAAS (XTXM202203), Elite Rice Plan of GDRRI (2022YG01), Guangdong Key Laboratory of New Technology in Rice Breeding (2020B1212060047), Seed Industry Revitalization Project of Special Fund for Rural Revitalization Strategy in Guangdong Province (2022NJS00004) and Special Foundation for Introduction of Scientific Talents of GDAAS (R2021YJ-XD001).

## Abbreviations

(DRS): Direct RNA sequencing
(lncRNA): long non-coding RNA
(snRNA): small nuclear RNA
(snoRNA): small nucleolar RNA
(PTC): peptidyl transferase center
(U): uridine
(RPUSD4): RNA Pseudouridine Synthase D4
(FCS1): Leaf Curly and Small 1
(Pus1p): Pseudouridine Synthase 1p
(PUS): pseudouridine synthase
(TruA): tRNA pseudouridine(38-40) synthase
(TruB): tRNA pseudouridine(55) synthase
(TruD): tRNA pseudouridine(13) synthase
(RluA): bifunctional tRNA pseudouridine(32) synthase/23S rRNA pseudouridine(746) synthase
(RNPs): ribonucleoproteins
(MS): mass spectrometry
(NGS): next-generation sequencing
(LC-MS/MS): liquid chromatography-tandem mass spectrometry
(2D-TLC): two-dimensional thin-layer chromatography
(ONT): Oxford Nanopore Technologies
(LC-MS): liquid chromatography-mass spectrometry
(GWAS): genome-wide association study
(sORF): small open reading frame
(eQTL): expression quantitative trait locus
(rG4): G-quadruplex
(RASP): RNA Atlas of Structure Probing
(CNN): convolutional neural network

