## Supplementary material for "Ψ-Atlas: an integrated atlas for pseudouridine epitranscriptome": Table 1

**Table 1. Comparison with other integrated Ψ modification related databases**

| Name | Ψ-Atlas | RMDisease V2.0 | EPITOMY | DirectRMDB | RMBase v3.0 | PRMD |
| --- | --- | --- | --- | --- | --- | --- |
| Data resources | | | | | | |
| Species | 55 | 4 | 2 | 25 | 50 | 20 |
| Experimentally validated sites | Yes | No | Yes | No | Yes | No |
| Next Generation Sequencing (NGS) | Yes | Yes | Yes | No | Yes | Yes |
| Direct RNA Sequencing (NanoPore) | Yes | No | No | Yes | No | Yes |
| Other specific regulation datasets | | | | | | |
| RBP binding sites | Yes | Yes | No | Yes | Yes | Yes |
| RNA Structures | Yes | Yes | No | No | Yes | Yes |
| micro-RNA target sites | Yes | Yes | No | Yes | Yes | Yes |
| Single nucleotide polymorphisms (SNPs) | Yes | Yes | No | Yes | Yes | Yes |
| Expression Quantitative Trait Loci (eQTLs) | Yes | No | No | No | No | Yes |
| GWAS sites | Yes | Yes | No | No | No | Yes |
| Small open reading frames (sORFs) | Yes | No | No | No | No | Yes |
| Alternative cleavage and polyadenylation (APA) sites | Yes | No | No | No | No | Yes |
| Tools and Visualization | | | | | | |
| Pseudouridine predictor | Yes | Yes | No | No | No | No |
| Variation effects on modifications | Yes | No | No | No | No | Yes |
| JBrowse | Yes | Yes | No | Yes | Yes | Yes |
| The ways to query datasets | | | | | | |
| Gene ID | Yes | Yes | Yes | No | No | Yes |
| Gene Name | Yes | Yes | Yes | Yes | No | Yes |
| Job ID | Yes | No | No | No | No | Yes |
| PubMed ID | Yes | No | Yes | No | No | Yes |
| Links | https://rnainformatics.org.cn/PsiAtlas | http://www.rnamd.org/rmdisease2 | https://epitomy.luddy.indianapolis.iu.edu | http://www.rnamd.org/directRMDB | https://rna.sysu.edu.cn/rmbase3/ | http://bioinformatics.sc.cn/PRMD |
